# Understanding the population structure of the GHQ-12: evidence for multidimensionality using Bayesian and Exploratory Structural Equation Modelling from a large-scale UK population survey

**DOI:** 10.1101/584169

**Authors:** Gareth J Griffith, Kelvyn Jones

## Abstract

Mental health and its complexity, measurement and social determinants are increasingly important avenues of research for social scientists. Quantitative social science commonly investigates mental health as captured by population screening metrics. One of the most common of these metrics is the 12-Item General Health Questionnaire (GHQ-12). Despite its canonical use as an outcome of interest in social science, the traditional use of the summed scores of summed questionnaires carries empirical and substantive assumptions which are often not fully considered or justified in the research. We outline the implications of these assumptions and the restrictions imposed by traditional modelling techniques and advocate for a more nuanced approach to population mental health inference. We use novel Exploratory Structural Equation Modelling (ESEM) on a large, representative UK sample taken from the first wave of the Understanding Society Survey, totalling 40,452 respondents. We use this to exemplify the potential of traditional, restrictive assumptions to bias conclusions and policy recommendations. ESEM analysis identifies a 4-factor structure for the GHQ-12, including a newly proposed “Emotional Coping” dimension. This structure is then tested against leading proposed factor structures from the literature and is demonstrated to perform better across all metrics, under both Maximum Likelihood and Bayesian estimation. Moreover, the proposed factors are more substantively dissimilar than those retrieved from previous literature. The results highlight the inferential limitations of using simple summed scores for mental health measurement. Use of the highlighted methods in combination with population studies offers quantitative social scientists the opportunity to explore predictors and patterns of underlying processes of population mental health outcomes, explicitly addressing the complexity and measurement error inherent to mental health analysis.

## 1 Introduction

Mental health measurement, specifically measurement of mood disorders, is notoriously difficult. This difficulty is largely due to the challenges of quantifying an individual’s mental health status robustly and the presence of ambiguity around tangible thresholds. As a result it is likely that there is substantial under-diagnosis of mental health disorders with as many as 74-85% of depressed individuals never receiving diagnoses of depression (Lecrubier, 2007; Verheij, 1996). It follows that the most under-reported group are individuals with ‘milder’ cases of depression (Garrard et al., 1998). As clinical outcomes under-report psychological morbidity, quantitative social scientists commonly address this by estimating psychological morbidity from questionnaires distributed to a sample of the population.

The 12-Item General Health Questionnaire (GHQ-12) is one of the most widely used responses in quantitative social science and epidemiology for the analysis of mental health trends. It constitutes 12 questions, each with 4 Likert response categories, which when answered are conventionally summed to give a single overall score lying on a notionally meaningful singular dimension (Goldberg and Hillier, 1979). The popularity of the GHQ-12 is mostly due to its ease of use, and capacity to reliably reproduce “remarkably robust” results contrasted with longer initial versions (Goldberg et al., 1997). Initially focusing on diagnostic purposes for specifically at-risk individuals, the GHQ-12 has since been translated and validated across multiple languages and countries as a screening tool for depression and depressive symptoms across populations (Creed and Evans, 2002; Hankins, 2008a; Pevalin, 2000; Smith et al., 2010). The UK is no exception to this, where the GHQ-12 has become the canonical mental health measure for UK-based population studies (Propper et al., 2005; Thomson and Katikireddi, 2018; Weich et al., 2003), largely due to its inclusion as principal mental health outcome for a series of popular large-scale surveys including the UK Household Longitudinal Survey (McFall, 2011) and the Health Survey for England (Mindell et al., 2012).

Despite being extensively validated using other leading mental health metrics, as well as longer versions of the same GHQ metric, there is still considerable debate as to what the GHQ-12 is truly capturing (Werneke et al., 2000; Ye, 2009). Due to its repurposed nature, different disciplines treat the GHQ-12 differently, seeking either to identify cases of ill-health (Goldberg and Bridges, 1987), or understand population trends across mental health spectra (Hu et al., 2007; Weich et al., 2003). This has led to numerous multidimensional factor structures being proposed in the literature, with little consensus on whether more complex structures are truly adding value or if they result simply from over-interpretation of substantively meaningless stochastic variation (Aguado et al., 2012; Hankins, 2008a).

The critical importance of understanding composite and complex measures such as the GHQ-12 has been clearly evidenced in recent genetic and psychological research. For instance, more strictly defined cases of depression produce higher estimates of genetic heritability for major depressive disorder (Cai et al., 2018); and mental illness and wellbeing, often used interchangeably in policy, can have widely differing covariates (Patalay and Fitzsimons, 2016; Westerhof and Keyes, 2010). Greater nuance in understanding has also been explicitly advocated by the UK Chief Medical Officer, who called for greater understanding of both positive and negative components of mental health (Davies, 2013, 2018).

This paper uses novel Exploratory Structural Equation Modelling techniques to estimate underpinning processes governing the GHQ-12 in the UK. This allows the relaxation of previously necessary methodological constraints which have been demonstrated to bias results towards simpler interpretations (Marsh et al., 2014). The factor structure provided by this procedure is subsequently evaluated against leading interpretations of GHQ-12 dimensionality. Moreover, all models are estimated using both Maximum Likelihood (ML) and Bayesian procedures allowing for more flexible estimation of non-normal values, giving less biased results for potentially skewed factor variances (Muthén and Asparouhov, 2010)

The rest of the paper is organised as follows. Firstly, a brief overview of the methodological assumptions underpinning traditional factor analysis methods is provided. We then critically review previous structures found in the literature for the GHQ-12, identifying common dimensions found in studies across sub-populations. We then consider the large-scale data to be analysed from Understanding Society and briefly outline a recently developed analytical approach, ESEM, which allows a less constrained approach to instrument decomposition than traditional factor analysis, with the potential to reveal a more nuanced structure. This analytical approach is then applied to identify an optimal factor structure for the study population, which is then compared with structures proposed and validated by previous studies. An extended discussion section concludes by exploring the implications and limitations of this newly found structure, with recommendations for best practice for quantitative social scientists in generating meaningful insight from complex mental health outcomes.

### 1.1 Understanding Mental Health Measurement

The desire for population screening metrics can be seen as symptomatic of a wider cultural shift in mental health perception away from an “absence of illness” perspective. Where early studies of the ecology of mental illness focused on diagnoses of psychosis (e.g. Faris and Dunham, 1939), evidencing change via reduction in illness, contemporary approaches advocate a more holistic approach aiming to measure improvements in “wellbeing” (World Health Organization, 2013). This necessarily involves the conceptualisation and characterisation of a mental health “spectrum”, beyond binary caseness. Whilst complexity is increasingly recognised as an intrinsic part of mental health measures (Gnambs and Staufenbiel, 2018; Hu et al., 2007), quantitative social science commonly takes the reductive view of reducing questionnaire responses to binary “cases” or as unidimensional constructs without adequate consideration of the methodological implications of doing so.

The underpinning theoretical assumption of using summed GHQ-12 scores is that mental health lies on a single spectrum, and that all variation in the outcome of interest occurs along that spectrum. This is referred to as unidimensionality and is particularly pertinent in literature which recommends or posits thresholds for the GHQ-12 above which an individual is considered at risk, or an “ill case” (Baksheev et al., 2011; Goldberg et al., 1998; Tait et al., 2003). In this literature, for caseness to be meaningful at the population level it is particularly critical that a unit increase can be assumed to be capturing the same change in mental status across individuals, an assumption that necessarily posits substantive equivalence in unit response change *between* items (unidimensionality), and *within* items (additivity). Ultimately all research treating the GHQ-12 as unidimensional, continuous or binary, necessarily requires the belief that a unit-increase in summed GHQ-12 score implies the same change wherever it occurs across the metric (Brodersen et al., 2007; Marsh and Bailey, 1991).

Moving beyond internal validity, the context-sensitivity of the interpretation of the GHQ-12 has been evident for some time (Werneke et al., 2000). Goldberg et al. (1998) noted the context-specific nature of thresholds in the GHQ-12, documenting the clear differences in specificity and discriminant capacity between-items across-countries. Despite this early acknowledgement, the problematic issue has been largely disregarded in subsequent literature due to necessary impositions of overly-simplistic analyses and validation measures.

Numerous studies have attempted to characterise the internal consistency of the GHQ-12 using relatively simple tests of Cronbach’s Alpha (Cronbach, 1951) or by conducting Confirmatory Factor Analysis (CFA) (Jöreskog, 1969). Of the two, CFA is considered the superior measure, providing robust evaluations of capacity for model replication (Dunn et al., 2014). However, it still requires a refinement of dimensionality to “simple structure”, implying zero cross-loadings across items (Asparouhov and Muthén, 2009a; Marsh et al., 2014). This means that each constituent item can only be empirically related to one underpinning construct, which has in turn been argued to lead to a reliance on overfitting models via model modification indices (MacCallum et al., 1992), this renders the ostensibly *confirmatory* analysis theoretically *exploratory* (Fabrigar et al., 1999; Schmitt, 2011). The issues surrounding this approach to social science metrics are not solely empirical, as there are also substantive concerns when using traditional CFA approaches (Conway and Huffcutt, 2003). The most pressing of these concerns is the imposition of zero cross-loadings, which are particularly unrealistic for measures such as the GHQ-12, as “nonzero cross-loadings are inherent in psychological measurement” (Marsh *et al.*, 2014, pp.88).

### 1.2 Proposed Factor Structures

Whilst longer versions of the GHQ are commonly accepted to be multidimensional (Graetz, 1991; Martin, 1999), consensus is less forthcoming for the GHQ-12. Of the variety of proposed interpretive structures suggested for the GHQ-12, there are three common interpretations. The most simplistic involves interpreting the GHQ-12 score as a unidimensional construct, taking the summed scores as a response, occasionally with an adjustment for positive and negative phrasing. This approach is backed up by a large body of research, which uses CFA to conclude that the unidimensional interpretation of the GHQ-12 is the most compelling (e.g. Aguado et al., 2012; French and Tait, 2004; Winefield et al., 1989). Despite this research concluding in support of unidimensional interpretation there is another consideration of importance here.

Studies commonly cite high correlations between modelled multidimensional factors as justification for simplifying to unidimensional interpretation (e.g. Gao *et al.*, 2004; Gouveia *et al.*, 2010; Padrón *et al.*, 2012; Fernandes and Vasconcelos-Raposo, 2013; Romppel *et al.*, 2013). Recent work using simulated data has called this justification into question, demonstrating this imposition of simple structure to artificially inflate correlations between modelled factors (Asparouhov and Muthén, 2009a; Marsh et al., 2014). Thus whilst it is not uncommon for reported correlations to be greater than 0.9 (e.g. Aguado et al., 2012; Campbell and Knowles, 2007) or 0.95 (e.g. Sweeting et al., 2009; Wang and Lin, 2011), taking these high correlations as justification for unidimensionality risks a self-fulfilling prophecy of simplicity begetting simplicity.

Whilst many studies call for unidimensional interpretation there are also numerous proposed multidimensional GHQ-12 structures. Several two-factor solutions have been proposed and validated using CFA techniques. These two factors most commonly involve a “Depression/Anxiety” construct and a “Social Dysfunction” construct (Andrich and Schoubroeck, 1989) (given by GHQ-12 items 2, 5, 6, 9, 10 and 11, and 1, 3, 4, 7, 8 and 12 respectively - see supplementary materials for full item list with response coding). “Depression/Anxiety” relates to the emotional component of psychological distress, whereas “Social Dysfunction” relates to the social functioning component of the individual experiencing the distress. This structure, albeit with different labelling, has at various points been identified as the best fit in data from the UK (Smith et al. 2010), New Zealand (Kalliath et al., 2004), Brazil (Gouveia et al., 2010), Japan (Suzuki et al., 2011), Germany (Schmitz et al., 1999), Italy (Politi et al., 1994) and Turkey (Kiliç et al., 1997). Systematic reviews and meta analyses also consistently identify these two factors most commonly in both two- and three-factor solutions (Gnambs and Staufenbiel, 2018; Picardi et al., 2001; Werneke et al., 2000). It is important to note that these groupings also align with the positive or negative phrasing of the constituent items. “Social Dysfunction” items are all positively worded and “Depression/Anxiety” items are all negatively worded, and as such there has been debate as to whether this structure simply reflects differences in phrasing (Hankins, 2008b).

Increasing the multidimensional complexity brings consideration of three-factor solutions, which most commonly identify “Social Dysfunction” and “Depression/Anxiety” constructs alongside a third construct referring to some variant of “Loss of Confidence”. This structure was initially identified by Worsley and Gribbin (1977) in the first factor analysis of the GHQ-12. They found the “Loss of Confidence” construct alongside “Social Performance” and “Anhedonia-Sleep Disturbance” (the latter of which approximate the Social Dysfunction and Depression/Anxiety factors respectively). For Worsley and Gribbin this third posited dimension was constructed of four items (6, 9, 10 and 11) all pertaining to feeling worthless or depressed, or losing confidence, evidence of which has been supported by subsequent work (Campbell et al., 2003; Penninkilampi-Kerola et al., 2006; Vanheule and Bogaerts, 2005). Notably, Worsley and Gribbin did not subsequently refine to a simple structure solution, as there were no other structures to test against, so this solution contains non-zero cross loadings.

A similar structure was published by Graetz in 1991. Graetz analysed a large scale (N= 8998) Australian sample, resulting in a three-factor structure comprising “Anxiety/Depression”, “Social Dysfunction” and “Loss of Confidence”. The Loss of Confidence factor is similar to that of Worsley and Gribbin – but is presented in the simple structure format of CFA, with non-zero cross-loadings eliminated. Graetz’s Loss of Confidence factor only loads on items 10 and 11, which are the only two of Worsley and Gribbin’s factor that do not also load on another factor. This 3-factor structure has been the most widely accepted multi-dimensional structure since its introduction, supported by subsequent work both in the UK (e.g. Cheung, 2002; Martin 7& Newell, 2005; Shevlin & Adamson, 2005) as well as elsewhere (Gnambs and Staufenbiel, 2018; Padrón et al., 2012).

Whilst several proposed structures from the literature are strongly validated, there are sampling issues with their initial design. Firstly, with the exception of the Graetz 3-factor model (Graetz, 1991), structures are commonly generated and subsequently validated using data with a small sample size and limited contextual scope. It is not uncommon for validation studies to report sample sizes under 500 (e.g. Khan et al., 2013; Martin and Newell, 2005). Moreover, studies with greater statistical power commonly draw samples from heavily context-specific populations such as primary care users (e.g. Werneke *et al.*, 2000) or high-school students (e.g. Suzuki *et al.*, 2011), and not the general population. Secondly, there is little methodological agreement on effective measures and fit criteria for recommending any given structure over another, despite efforts to establish consistent approaches (Conway and Huffcutt, 2003; Hu and Bentler, 1999; Marsh et al., 2004).

There is clear substantive interest to both social and health researchers in moving beyond understanding the patterning of observed, aggregated measures of complex phenomena such as mental health. It is of critical importance to develop a better understanding of the underpinning processes driving these measures. Empirically, this means partitioning variance in response to each item into common and unique variation across questions, thus factoring for individual variability in response (Conway and Huffcutt, 2003). There is clearly a wealth of longitudinal data available for the GHQ-12 in the UK, collected alongside rich demographic and spatial data in several panel studies. In light of newer methodological developments in the modelling of complex responses, this wealth of data needs comprehensively re-evaluating to ensure it is fully exploited. This study proposes a multidimensional interpretation of latent factors patterning GHQ-12 responses, allowing insight into underpinning processes rather than manifest outcomes. It further aims to clarify and make explicit the empirical and substantive benefits of doing so, in using novel methodological developments to consider more complex dimensionality, in the quantitative analysis of a subject as nuanced and individually heterogeneous as mental health.

## 2 Data and Method

### 2.1 Data

This study uses the first wave of the large scale Understanding Society Survey (US), a nationally representative annual panel survey of over 43,000 UK respondents from over 26,000 households (McFall, 2011). Data collection for Wave 1 took place between January 2009 and March 2011. The GHQ-12 was administered via face-to-face interview and completed by 40,452 respondents. For a comprehensive overview of Understanding Society Data Collection and methods see Quality Profile (Lynn and Knies, 2016). Table 1 gives the response characteristics of the survey sample. For initial exploration sample weights were taken and applied from Wave 1. Sample weights were applied to all but Confirmatory Bayesian Analyses, as weights are not supported in Bayesian estimation.

**Table 1:**
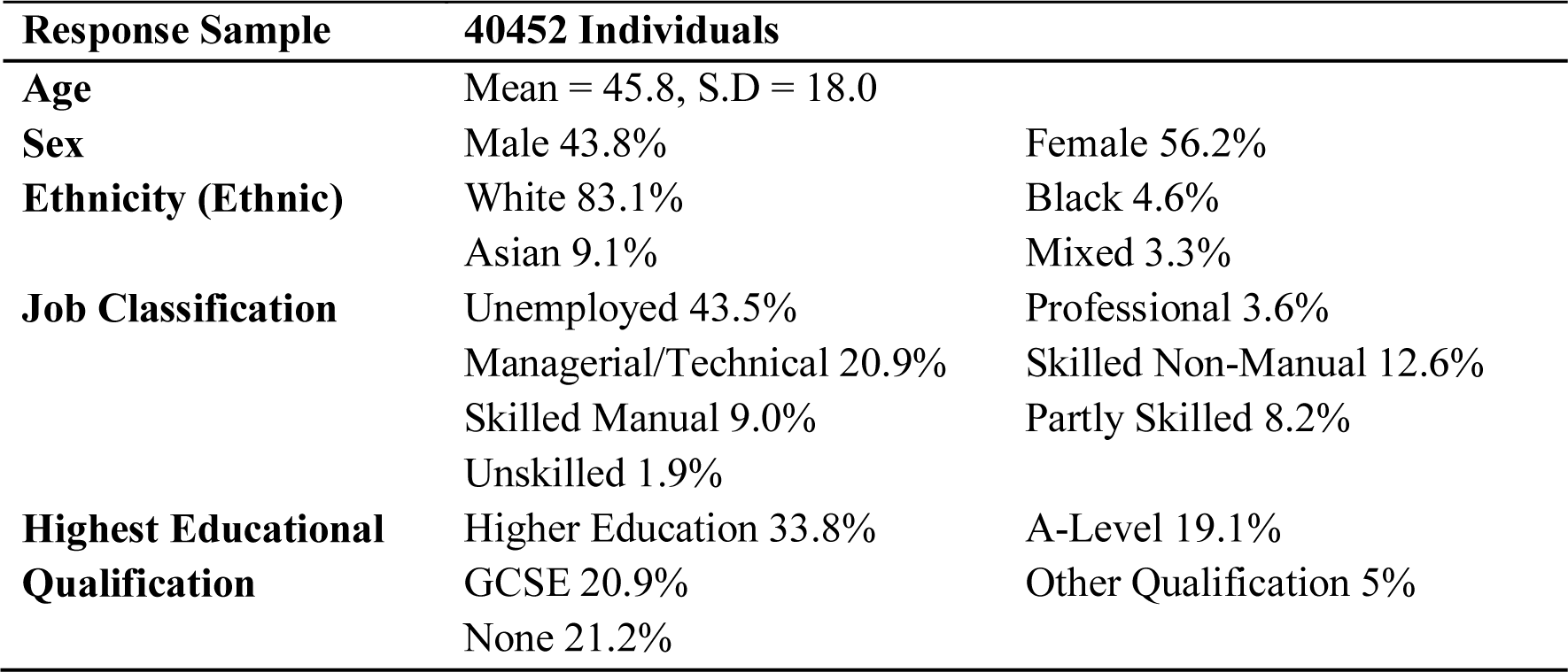
Response Characteristics of Survey Sample completing the GHQ-12 (Understanding Society Wave 1)

There is strong evidence of different response patterning across positive and negative questions as part of the GHQ-12, shown in Figure 1. It can be seen that there are strongly different modal responses for positive and negative items, giving rise to the common factor structures reflecting positive and negative items. Despite strong patterning in responses which is somewhat corrected for in binary interpretation, Likert scores have been demonstrated to allow greater discrimination between different dimensional structures (Campbell and Knowles, 2007; Smith et al., 2013). It is for this reason that we use full information Likert scoring here. Items are all coded such that higher scores indicate greater distress.

**Figure 1:**
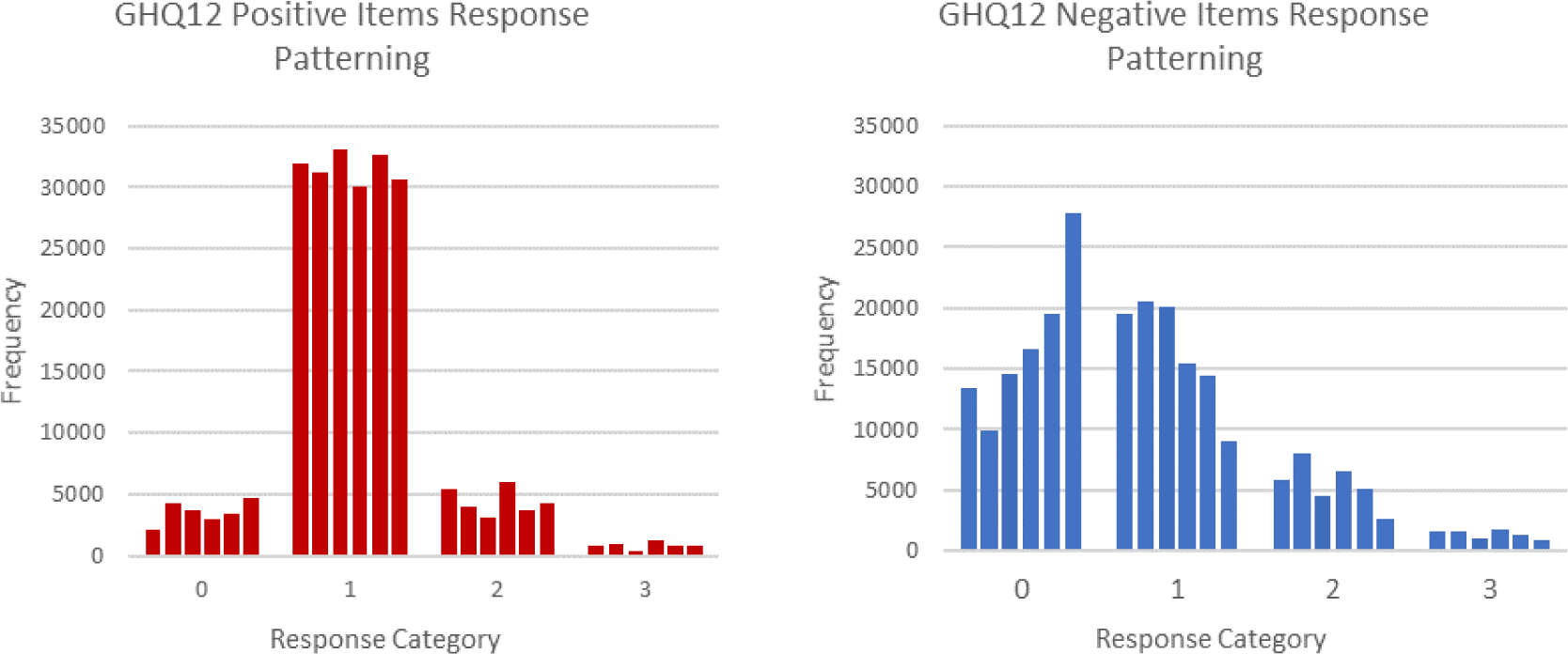
Graph illustrating the differential frequency of response patterning of positively (1,3,4,7,8,12) and negatively (2,5,6,9,10,11) phrased items in the GHQ-12 from 2009 Wave of Understanding Society.

### 2.2 Methodology

To explore the GHQ-12 data a novel approach termed Exploratory Structural Equation Modelling (ESEM) is deployed (Asparouhov and Muthén, 2009). ESEM represents a combination of the best elements of restrictive CFA and unstructured exploratory factor analysis (EFA). The key contribution of ESEM methodology for this research is the specification of non-zero cross-loadings on constituent items (Asparouhov and Muthén, 2009). As outlined above, in a literature which commonly cites high factor correlations as justification for refining to unidimensional interpretation, it is especially important to guard against inflated estimates of factor correlations which result from imposing zero cross-loadings (Asparouhov et al., 2015; Marsh et al., 2014, 2010). Beyond the empirical, there is also a substantive argument for not refining to simple structure when analysing complex social outcomes such as psychological constructs. As multiple and inter-related underpinning processes are likely to give rise to any specific outcome most items will have multiple determinants and thus nonzero cross-loadings ought not to be viewed an aberration but a logically anticipated representation of complex, underpinning constructs (Marsh *et al.*, 2014). ESEM is carried out within the Mplus software environment (Muthén and Muthén, 1998-2018).

The model is estimated using a two-step procedure. Firstly, a series of EFA solutions in which each item loads on each and every of 2-5 constructs is estimated using Geomin rotation (ε = 0.01), the optimal rotation when little is known about the true underlying structure, and when suspected variable complexity is greater than one – that is, there is an expectation that there will be cross-loadings (Browne, 2001; McDonald, 2005). Loading retention is dictated by a simple cutoff of EFA loadings <0.15. As the need to exclude loadings purely for methodological reasons has been relaxed, the value of 0.15 is a far more tolerant exclusion criterion than typically advocated in the literature. To test this criterion, models were rerun with all loadings present but target rotation of 0 for these small EFA loadings. This resulted in all non-target loadings having absolute values below 0.1, thus they were omitted in the confirmatory component of the model comparison.

The Likert GHQ-12 responses are specified as ordinal variables and ML estimation of confirmatory analyses was carried out using the mean and variance-adjusted weighted least squares estimator as this presents the best method for categorical or ordered data (Schmitt, 2011). This estimation specifies a probit regression equation for each item on the related factor (Muthén & Muthén, 1998-2011, pp. 62). These can be interpreted as likelihood changes in the log-odds of changing response category on the response variable, illustrating the strength of the relation between the underpinning dimension and the probability of response change in the associated item (Chen, 2007).

The models were all repeated using Bayesian estimation, allowing for less biased estimation of non-normal response variances than common ML methods (Muthén and Asparouhov, 2010). The distributions of estimates are highly likely to be non-normal due to the constrained nature of variances and correlations, with variances having a lower limit of zero, and correlations bounded by 1 and −1. Bayesian estimation does not require the ML assumption of normality of the parameter estimates, with prior variance-covariance estimates instead drawn from an inverse-Wishart distribution (Asparouhov and Muthén, 2010).

Model fit for the ML estimation was evaluated using the Comparative Fit Index (CFI), Tucker-Lewis Index (TLI), Root Mean Square Error of Approximation (RMSEA) and the Weighted Root Mean Square Residual (WRMR). Evidence of good fit was taken from guidelines proposed by Bentler (2007) and Muthén and Muthén (2014). For the Bayesian models, fit is evaluated using Posterior Predictive Checking (PPC) for which we present posterior predictive p-values and 95% credible intervals for each model. In Mplus PPC is an extension of the likelihood ratio statistic taken to be an indication of the model’s capacity to reproduce the data and summarise the posterior distribution of the residuals (Asparouhov and Muthén, 2010). As such, the statistic is upward biased by sample size so we consider evidence of improvement in predictive capacity as reduction towards zero of the predictive credible interval (Gelman et al., 1996; Marsh et al., 2004). We also present mean absolute factor correlation values as evidence of substantive dissimilarity between modelled constructs with lower values indicating greater discriminant validity.

## 3 Results

Initial analysis specified a series of EFAs with between 2 and 5 factors, the results of which are given in Table 2. In the traditional ML estimation, goodness of fit statistics evidence better model improvement for each added dimension up to a five-factor solution. In the Bayesian estimation, it is more explicit that none of the specifications provide an adequate fit to reproduce the 40,452 individuals’ mental health response, as shown by the consistent posterior p-values of 0.000. However, there is evidence of improvement of fit in the posterior credible interval. Predictive capacity improves with added factors up to the fourth factor; however, the five-factor solution is less robustly estimated. Reduction in the lower bound shows that a fifth factor could serve to improve predictive capacity but the increased upper bound suggests it is more probable that it will reduce the predictive capacity of the model. Therefore, it is the four-factor structure that is carried forward as it offers the most robust reproduction of observed values.

**Table 2:**
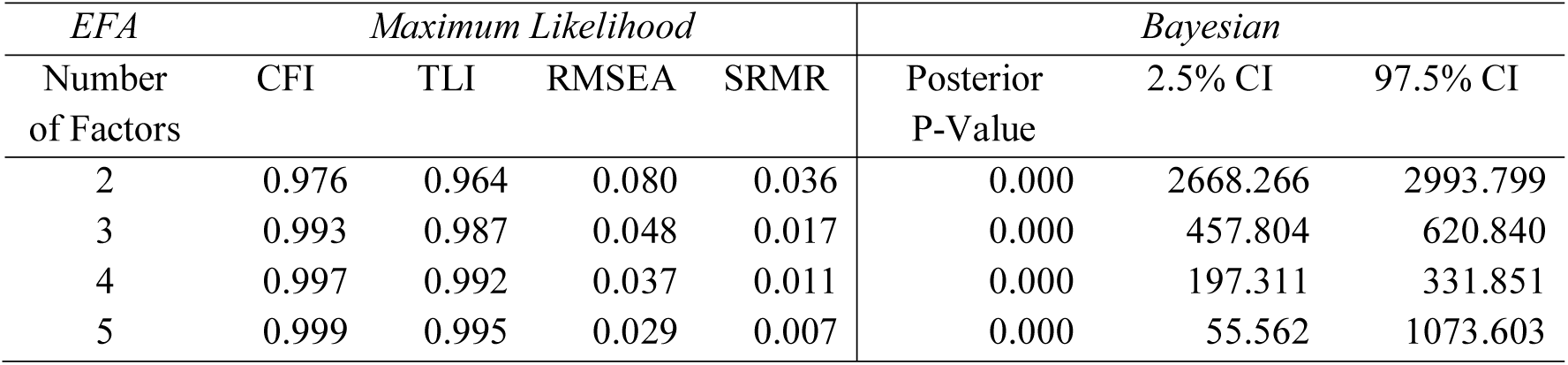
Bayesian and ML Fit statistics for EFA factor solutions for the GHQ-12 with 2, 3, 4 and 5 factors.

Having identified that the four-factor solution was the most appropriate from the estimation of the initial EFA whilst all cross-loadings are specified and estimated, the subsequent solution needed refining to a testable structure. This was implemented by re-specifying the model with all cross-loading values under 0.15 being omitted. The loadings can be interpreted as probit regression coefficients of each item on the unit-standardised response factor and are directly comparable both within and between factors.

Table 3 gives the loadings for the four-factor ESEM solution. The four factors are labelled “Lowered Self Worth”, “Social Dysfunction”, “Stress” and “Emotional Coping”. These were chosen in accordance with existing names for previously identified factors from the literature (Graetz, 1991; Martin, 1999). The main difference from previous structures in the literature is that items 1, 2, 4, 5, 6, 7, 9 and 12 now load on multiple underlying dimensions. The loadings of Factors 1 and 2 are consistent with previous literature, and thus have been given similar names to reflect this. However, they differ empirically because they contain cross-loading items, but the broad differentiation between the positive and negatively worded items holds true. Higher individual factor scores across Factors 1 and 2 indicate higher levels of mental distress.

**Table 3:**
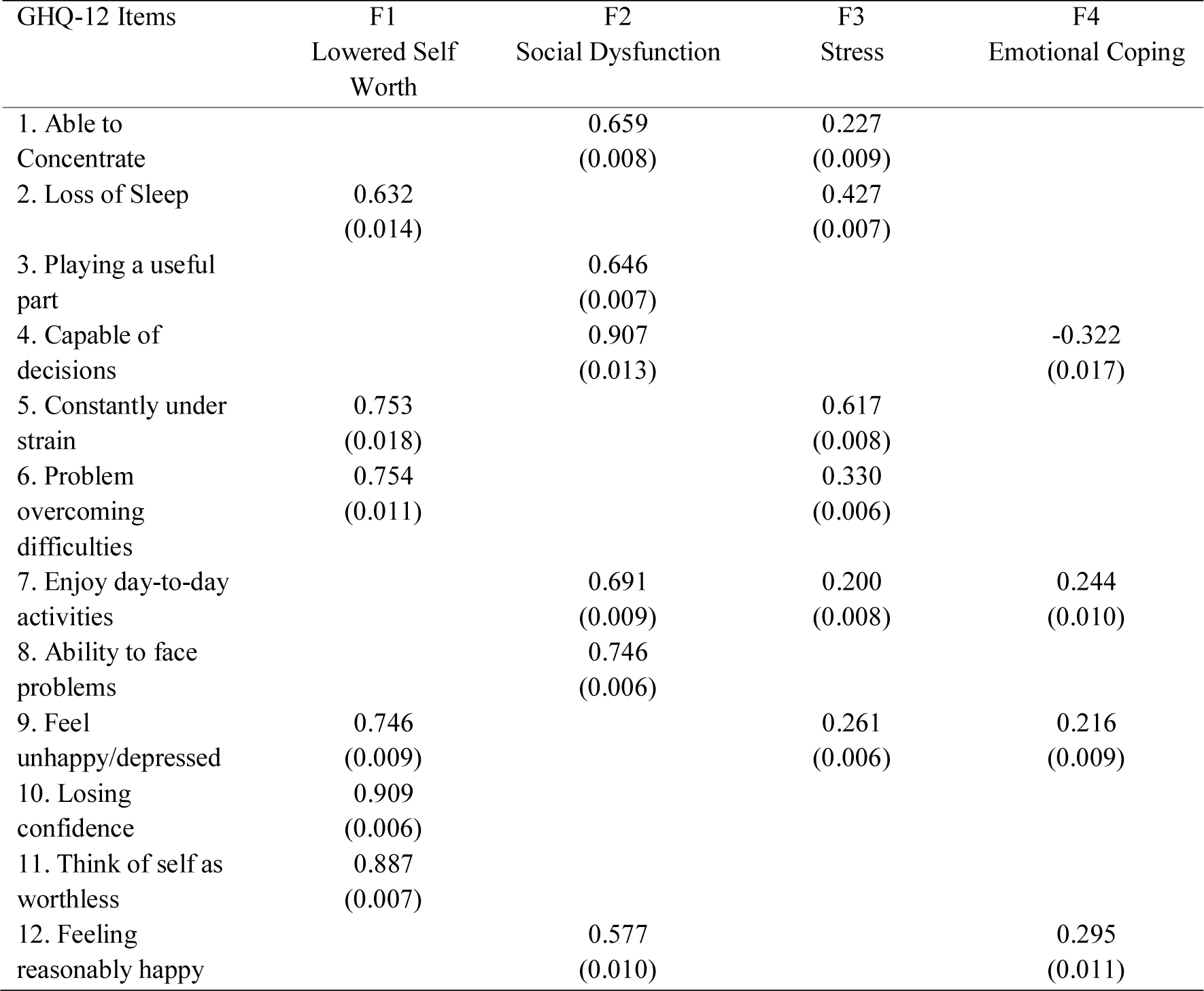
Standardised Factor Loadings for 4-Factor ESEM Model of the GHQ-12 using Bayesian estimation. Standard Errors in parentheses.

| GHQ-12 Items | F1<br>Lowered Self<br>Worth | F2<br>Social Dysfunction | F3<br>Stress | F4<br>Emotional Coping |
| --- | --- | --- | --- | --- |
| 1. Able to<br>Concentrate |  | 0.659<br>(0.008) | 0.227<br>(0.009) |  |
| 2. Loss of Sleep | 0.632<br>(0.014) |  | 0.427<br>(0.007) |  |
| 3. Playing a useful<br>part |  | 0.646<br>(0.007) |  |  |
| 4. Capable of<br>decisions |  | 0.907<br>(0.013) |  | -0.322<br>(0.017) |
| 5. Constantly under<br>strain | 0.753<br>(0.018) |  | 0.617<br>(0.008) |  |
| 6. Problem<br>overcoming<br>difficulties | 0.754<br>(0.011) |  | 0.330<br>(0.006) |  |
| 7. Enjoy day-to-day<br>activities |  | 0.691<br>(0.009) | 0.200<br>(0.008) | 0.244<br>(0.010) |
| 8. Ability to face<br>problems |  | 0.746<br>(0.006) |  |  |
| 9. Feel<br>unhappy/depressed | 0.746<br>(0.009) |  | 0.261<br>(0.006) | 0.216<br>(0.009) |
| 10. Losing<br>confidence | 0.909<br>(0.006) |  |  |  |
| 11. Think of self as<br>worthless | 0.887<br>(0.007) |  |  |  |
| 12. Feeling<br>reasonably happy |  | 0.577<br>(0.010) |  | 0.295<br>(0.011) |

The third factor, here termed “Stress” is seen to load most strongly on items associated with feeling under strain and loss of sleep. It is very similar to a factor found in one of the earliest factor analyses of the GHQ-12 by Worsley and Gribbin (1977), differing only in that they found it also loaded on item 12.

The emergence of the fourth factor, termed “Emotional Coping” is a distinctive finding. It is most notable for having both positive and negative loadings. It is *negatively* associated with Item 4 – “feeling capable of making decisions”, but *positively* associated with feeling unhappy or depressed, not enjoying day-to-day activities and not feeling happy. As such individuals with high scores on Emotional Coping are those who feel negative as captured by items 7, 9 and 12, whilst feeling capable of making decisions as captured by item 4, indicating a degree of (at least perceived) perseverance in the face of the distress. The negative loading highlights an ambiguity in interpreting latent variables. That is, the interpretation is imposed by the researcher, thus an empirically identical interpretation would be the inverse, with negative loadings on Items 7, 9 and 12 and a positive loading on Item 4. This would in turn invert the poles of the underlying coping dimension, with higher scores indicating positive rather than negative outcomes. Emotional Coping is structurally most similar to the “Sleep Disturbance/Anhedonia” construct found in work by Worsley and Gribbin (1977) although there is a key difference in the negative loading of Item 4, which is clearly associated with positive functioning.

Having detailed the theoretical implications of the proposed factor structure it is necessary to understand the substantive and empirical implications of the structure. To do this we evaluate the substantive dissimilarity between factor constructs as measured by the correlations between those constructs.

Table 4 shows the modelled factor correlations. The factors with the highest correlation are Lowered Self Worth and Social Dysfunction, with a coefficient of 0.68. Although it is the highest identified in this structure, it is still low relative to that found in the existing literature. Stress is the most statistically dissimilar and therefore most substantively distinct factor, exhibiting uniformly low correlations with the other constructs.

**Table 4:**
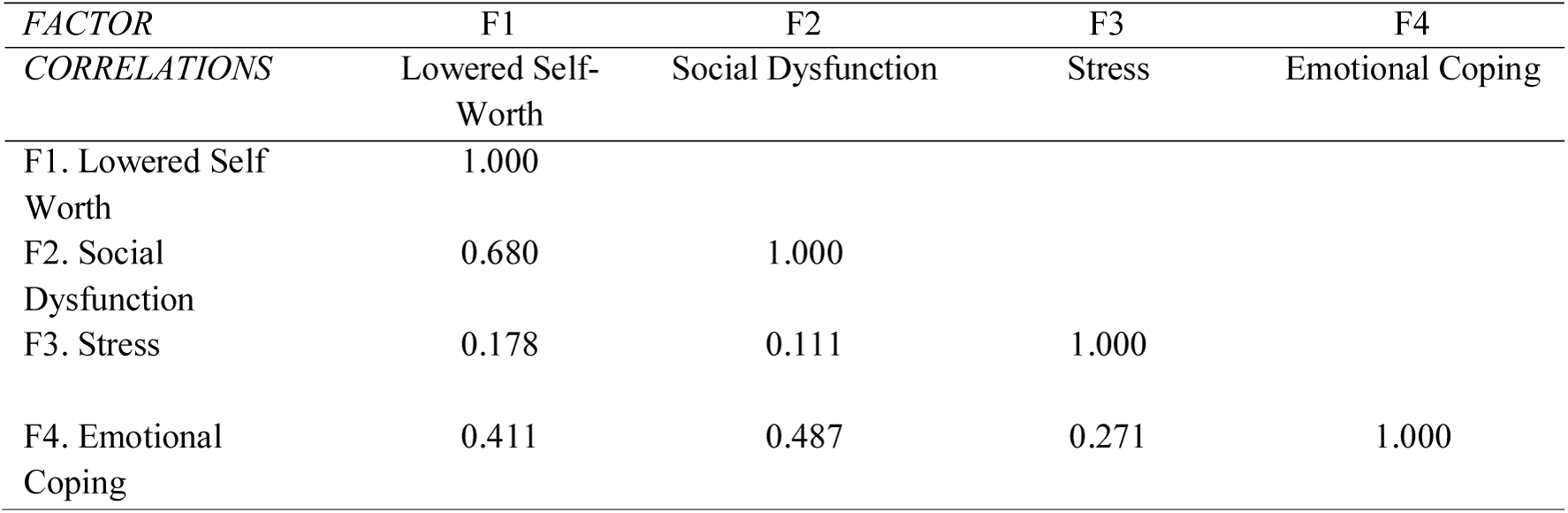
Modelled Factor Correlations from the Four Factor ESEM Solution for the GHQ-12

It should be noted that correlation coefficients overestimate the predictive capacity of each latent variable on another. The absolute proportion of variation in one variable that could be predicted solely from knowing the other is given by the squared value of the correlation coefficients (Kish, 1954). For instance, knowing the modelled Lowered-Self-Worth scores for all individuals would only allow the prediction of 46.24% (0.4624) of the variation in Social Dysfunction scores, despite these factors having the highest modelled correlation of 0.68. This is even more stark for the correlation between Stress and Social Dysfunction, with a true predictive capacity of 1.2%, leaving 98.8% of variation in one unexplained by the other.

Having identified a parsimonious summary of data from our model it is important to connect back to the wider literature on mental health structures. The literature has proposed many different structures, seen in Table 5, which gives context to seven exemplar proposed structures against which we evaluate the structure proposed here.

**Table 5:**
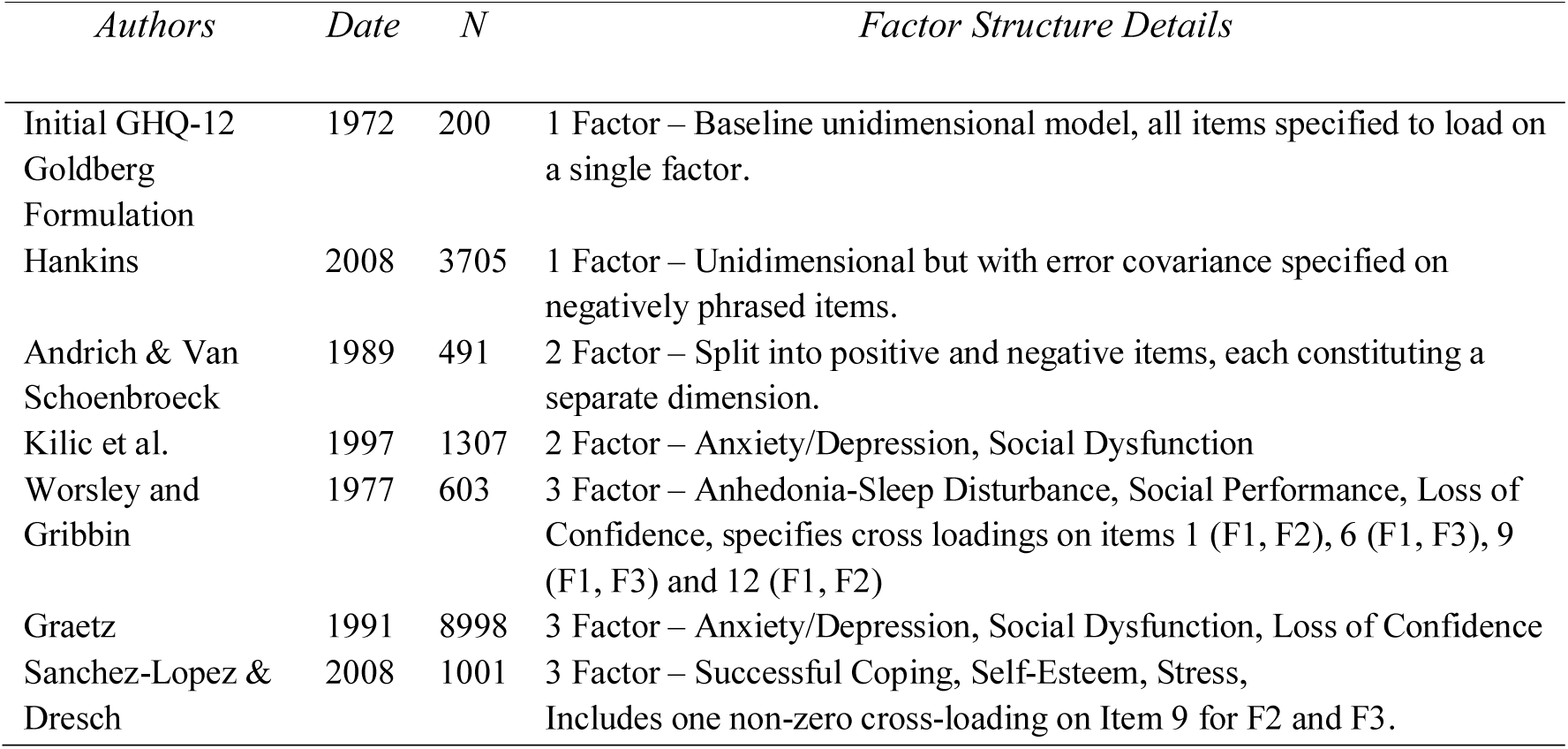
Seven exemplar studies of the range of factor analytical structures obtained from the GHQ-12 data. See Supplementary Materials for diagrammatic representation of structures.

| *Authors* | *Date* | *N* | *Factor Structure Details* |
| --- | --- | --- | --- |
| Initial GHQ-12<br>Goldberg<br>Formulation | 1972 | 200 | 1 Factor – Baseline unidimensional model, all items specified to load on a single factor. |
| Hankins | 2008 | 3705 | 1 Factor – Unidimensional but with error covariance specified on negatively phrased items. |
| Andrich & Van<br>Schoenbroeck | 1989 | 491 | 2 Factor – Split into positive and negative items, each constituting a separate dimension. |
| Kilic et al. | 1997 | 1307 | 2 Factor – Anxiety/Depression, Social Dysfunction |
| Worsley and<br>Gribbin | 1977 | 603 | 3 Factor – Anhedonia-Sleep Disturbance, Social Performance, Loss of Confidence, specifies cross loadings on items 1 (F1, F2), 6 (F1, F3), 9 (F1, F3) and 12 (F1, F2) |
| Graetz | 1991 | 8998 | 3 Factor – Anxiety/Depression, Social Dysfunction, Loss of Confidence |
| Sanchez-Lopez &<br>Dresch | 2008 | 1001 | 3 Factor – Successful Coping, Self-Esteem, Stress,<br>Includes one non-zero cross-loading on Item 9 for F2 and F3. |

**Table 6:** Bayesian and ML Fit Statistics for the 8 evaluated factor structure defined in Table 6.

| **Factor Structure** | No. of Factor s | WRMR | CFI | TLI | RMSE A | Posterior predictive P | Lower Bound | Upper Bound | Mean Absolute Factor Correlation |
| --- | --- | --- | --- | --- | --- | --- | --- | --- | --- |
| *Unidimensional* | 1 | 14.776 | 0.914 | 0.895 | 0.146 | 0.000 | 7123.436 | 7634.537 | - |
| *Hankins* | 1 | 6.808 | 0.981 | 0.967 | 0.082 | 0.000 | 1505.348 | 1764.743 | - |
| *Kilic et al.* | 2 | 12.826 | 0.934 | 0.918 | 0.129 | 0.000 | 5268.267 | 5709.397 | 0.822 |
| *Andrich & van Schoubroeck* | 2 | 9.018 | 0.967 | 0.959 | 0.091 | 0.000 | 3521.095 | 3892.429 | 0.777 |
| *Sanchez-Lopez & Dresch* | 3 | 8.246 | 0.972 | 0.963 | 0.087 | 0.000 | 2824.202 | 3152.636 | 0.772 |
| *Graetz* | 3 | 7.917 | 0.974 | 0.966 | 0.083 | 0.000 | 2459.754 | 2769.927 | 0.783 |
| *Worsley & Gribbin* | 3 | 5.505 | 0.986 | 0.981 | 0.063 | 0.000 | 1199.430 | 1430.393 | 0.665 |
| *ESEM 4Fac* | 4 | 3.247 | 0.995 | 0.992 | 0.039 | 0.000 | 376.571 | 526.539 | 0.356 |

Table 7 presents fit statistics for all seven models alongside the proposed ESEM solution. The mean absolute factor correlation is also presented for each specification to provide an estimate of the dissimilarity of the factors estimated in the model structure. The four-factor solution provides the best fit across every measure of fit under both traditional and Bayesian estimation.

The best performing structure outside of the ESEM solution is the original Worsley and Gribbin (1977) structure with its non-zero cross-loadings. Factor structures which address the inter-dependencies of the items via error covariance or non-zero cross-loadings perform well across all the fit statistics, which again is an argument for the adoption of a more realistically complex specification of mental health. The benefit of cross-loadings is most clearly borne out in the mean absolute factor correlation, which is far lower for the four-factor structure than other solutions.

## 4 Discussion

The results of this study show that of the factor structures tested here the four-factor structure provides the best description of GHQ-12 responses from the Understanding Society data, Wave 1. Additionally, this is evidenced not just by traditional fit statistics, but by the modelled factors being the most substantively dissimilar as evidenced by the mean absolute factor correlation. This involves the specification of two previously underexplored constructs, here termed “Stress” and “Emotional Coping”. Emotional Coping is particularly notable as the presence of a negative loading evidences the capacity for underpinning constructs to mask the presenting of psychological distress in the aggregated metric.

Within the wider GHQ-12 literature there is little mention of dimensions analogous to the Stress and Emotional Coping structures beyond Worsley and Gribbin’s early analysis (1977). Stress-related constructs are also proposed in a structure drawn from a Spanish population (Sánchez-López and Dresch, 2008), and notably the “thematic analysis” of GHQ-12 content by Martin (1999). These constructs would not have been discoverable using traditional CFA techniques as they consist largely of cross-loadings. The incorporation of these constructs considerably improved model fit, and the low modelled factor correlations seems to suggest that they are capturing substantively different processes. These low correlations in the results seem to support suggestions that high correlations may simply be an artefact of a restrictive modelling procedure and as such we caution refining to simpler structures based purely on apparently high factor correlations (Marsh et al., 2009).

The key message of this study is the capacity of large-scale datasets to contribute more comprehensive understandings of mental health outcomes in large, heterogeneous populations. Whilst this is not a new idea (Hu et al., 2007; Mukuria et al., 2014), the adoption of less stringent exclusion criteria in model selection and the incorporation of ESEM methodologies is something that is underexplored in large-scale survey analysis. Moreover, the statistical power afforded by these large-scale surveys using decomposed metrics has the potential to offer a more comprehensive understanding of the similarities between different underpinning processes across as well as within metrics.

That said, it is important to highlight the limitations of this study. Whilst the structure proposed here provides the best fit for the data in Wave 1 of Understanding Society in the UK (McFall, 2011), it is necessary to test on a wider range of data beyond this spatiotemporally specific dataset. More research is required in order to understand fully what can be gained from the modelling of decomposed processes underpinning survey instruments. Firstly, whilst sample weights were used in the generation of the factor structure, the structure is still specific to the current dataset. Another of the key contributions of the ESEM method is the capacity for comprehensive tests of measurement invariance over subgroups within a population (Marsh et al., 2009). Whilst it has been demonstrated that this structure provides a superior fit across the full population, the degree of measurement invariance for the structure should be investigated across different geographical and demographic groups. There is considerable literature validating the overall GHQ-12 as a screening instrument over time and space (e.g. Gnambs and Staufenbiel, 2018), there is far less written on the temporal or geographical stability of its underpinning latent structure (Goldberg et al., 1997). The structure should be further validated across spatio-temporal contexts using the wealth of existing GHQ-12 data.

Secondly, whilst it is clear that fit indices favour the proposed structure, it is important to establish that the proposed structure here offers more substantively in terms of predictive capacity than simpler structures. Recommendations for fit indices come from studies using sample sizes far smaller than those used here (Bentler, 2007; Hu and Bentler, 1999). As such, in isolation there are multiple structures tested here which would be accepted as “adequate” under such criteria. Whilst the Bayesian fit statistics presented are more robust to sample size, more work is needed on the appropriateness of fit-statistic thresholds in the face of increasingly large sample sizes (Muthén and Asparouhov, 2010). Proposed structures must be evaluated not just numerically, but on a theoretical basis, identifying whether the more complex structure offers greater understanding. The mean absolute factor correlations give some indication of this in terms of substantive dissimilarity between constructs. However comprehensive evaluation of this requires the investigation of the predictors of the different constructs, demographically, geographically and socially to evidence whether they truly add to our understanding of different processes. As such, further work is needed using the decomposed constructs as responses with data for which the structure is validated.

Similarly, given the use of the GHQ-12 as an external validation instrument (e.g. Mukuria *et al.*, 2014), it is important to incorporate these more nuanced understandings in subsequent studies evaluating what is being captured by other existing or new metrics (e.g. Tennant *et al.*, 2007). Adopting this approach will allow researchers to go further than modelling similarities in manifest responses to understanding similarities between underpinning mental health processes across populations. One such avenue is to investigate if any of the structures identified here are associated with similar underpinning processes of more recently developed well-being metrics such as the Short Warwick-Edinburgh Mental Well-Being Scale (Stewart-Brown et al., 2009). This will allow an empirical contribution to the debate on the contested relationship of mental illness and well-being (Westerhof and Keyes, 2010).

There are further limitations in the interpretation of the resulting constructs. It has been suggested in GHQ-12 literature that multidimensional factor structures are simply a product of over-interpretation of spurious variance in negatively worded items (Hankins, 2008a). It is important to note the possibility of this being the case in this data, the unidimensional correlated-error model performs very well, given its brevity. It initially seems reasonable to infer from this the potential for multidimensionality being solely the result of phrasing, but only if one assumes unidirectional causality, i.e. items were grouped into positive and negative items at random, rather than based upon conceptually different measurements (Gnambs and Staufenbiel, 2018). Empirically these two scenarios would present identically, although it seems reasonable to assume greater likelihood of the latter.

Replicating this analysis across different temporal and spatial contexts is a clear avenue of further research. Whilst understanding the stability of the structure across contexts is undoubtedly important, it is also important to demonstrate its worth in further developing understanding of what is being captured by these measures in large scale surveys. It is important, therefore, to evaluate the insight gained by decomposing complex metrics by taking them as responses in spatial, social, structural and genetic epidemiological studies. It is clearly of greater benefit to social scientists in any analysis positing putative causal mechanisms to robustly link predictors with *underpinning processes* of mental health than with aggregate, unidimensional questionnaire responses.

In conclusion this study, despite the above limitations, is one of the first to combine Bayesian and ESEM methodology with large-scale survey data in the UK and has demonstrated the inferential benefits of doing so for research into population-level mental health determinants. Further research is needed to validate these findings, using data from wider contexts, and contextualise them against differing mental health responses.

