## Supplementary Table 1 & 2 for "Understanding the population structure of the GHQ-12: evidence for multidimensionality using Bayesian and Exploratory Structural Equation Modelling from a large-scale UK population survey"

| - GHQ-12 Item | ***Scoring*** | | | |
| --- | --- | --- | --- | --- |
| **“Have you recently?”** | **0** | **1** | **2** | **3** |
| 1. Been able to Concentrate on what you’re doing? | Better than usual | Same as usual | Less than usual | Much less than usual |
| *2.* *Lost much sleep over worry?* | Not at all | No more than usual | Rather more than usual | Much more than usual |
| 3. Felt you were playing a useful part in things? | More than usual | Same as usual | Less so than usual | Much less than usual |
| 4. Felt capable of making decisions? | More so than usual | Same as usual | Less so than usual | Much less than usual |
| *5.* *Felt constantly under strain?* | Not at all | No more than usual | Rather more than usual | Much more than usual |
| *6*. *Felt you couldn’t overcome your difficulties?* | Not at all | No more than usual | Rather more than usual | Much more than usual |
| 7. Been able to enjoy normal day-to-day activities? | More than usual | Same as usual | Less so than usual | Much less than usual |
| 8. Been able to face up to your problems? | More so than usual | Same as usual | Less so than usual | Much less than usual |
| *9. Been feeling unhappy and depressed?* | Not at all | No more than usual | Rather more than usual | Much more than usual |
| *10. Been losing confidence in yourself?* | Not at all | No more than usual | Rather more than usual | Much more than usual |
| *11. Been thinking of yourself as a worthless person?* | Not at all | No more than usual | Rather more than usual | Much more than usual |
| 12. Been feeling reasonably happy, all things considered? | More than usual | Same as usual | Less so than usual | Much less than usual |

List of GHQ-12 questionnaire items with scoring for responses (Goldberg et al., 1997). Negatively phrased questions highlighted in italics.

| GHQ Item | Uni | Hankins (2008) | Kilic et al. (1997) | | Andrich & Van Schoubroeck (1989) | | Sanchez-Lopez & Dresch (2008) | | | Graetz (1991) | | | Worsley and Gribbin (1977) | | | ESEM (2019) | | | |
| --- | --- | --- | --- | --- | --- | --- | --- | --- | --- | --- | --- | --- | --- | --- | --- | --- | --- | --- | --- |
| Factor # | 1 | 1 | 1 | 2 | 1 | 2 | 1 | 2 | 3 | 1 | 2 | 3 | 1 | 2 | 3 | 1 | 2 | 3 | 4 |
| **1** | x | x |  | x | x |  | x |  |  | x |  |  | x | x |  | x |  | x |  |
| *2* | x | x! | x |  |  | x |  |  | x |  | x |  | x |  |  |  | x | x |  |
| **3** | x | x |  | x | x |  | x |  |  | x |  |  |  | x |  | x |  |  |  |
| **4** | x | x | x |  | x |  | x |  |  | x |  |  |  | x |  | x |  |  | x |
| *5* | x | x! | x |  |  | x |  |  | x |  | x |  | x |  |  |  | x | x |  |
| *6* | x | x! | x |  |  | x |  | x |  |  | x |  | x |  | x |  | x | x |  |
| **7** | x | x |  | x | x |  | x |  |  | x |  |  | x | x |  | x |  | x | x |
| **8** | x | x |  | x | x |  | x |  |  | x |  |  |  | x |  | x |  |  |  |
| *9* | x | x! | x |  |  | x |  | x | x |  | x |  | x |  | x |  | x | x | x |
| *10* | x | x! | x |  |  | x |  | x |  |  |  | x |  |  | x |  | x |  |  |
| *11* | x | x! | x |  |  | x |  | x |  |  |  | x |  |  | x |  | x |  |  |
| **12** | x | x |  | x | x |  | x |  |  | x |  |  | x | x |  | x |  |  | x |

Factor Structure of all eight tested GHQ-12 solutions including new ESEM solution. “x” indicates a specified loading, “!” indicates error covariance. Positively worded Items emboldened, Negatively worded items italicised.
